# Evaluation of RNA-seq normalization methods using challenging datasets

**DOI:** 10.1101/401679

**Authors:** Severin Uebbing

## Abstract

RNA-seq is a powerful tool for both discovery and experimentation. Most RNA-seq studies rely on library normalization to compare samples or to reliably estimate quantitative gene expression levels. Over the years a number of RNA-seq normalization methods have been proposed. Review studies testing these methods have provided evidence that commonly used methods perform well in simple normalization tasks, but their performance in challenging normalization tasks has yet to be evaluated. Here I test RNA-seq normalization methods using two challenging normalization scenarios. My assessment reveals surprising shortcomings of some commonly used methods and identifies an underappreciated method as the most promising normalization strategy for common, yet challenging RNA-seq experiments.

## Introduction

High-throughput RNA sequencing (RNA-seq) has proven a powerful tool for discovery and experimentation since its first introduction over a decade ago. It was realized early on that quantitative normalization of sequencing results is crucial for any downstream analysis [1]; normalization is used to handle differences in transcriptome composition, sequencing depth and other technical artifacts, for instance from combining data from different sources. Optimally, RNA-seq normalization methods should be able to deal with these issues and allow combining data from samples that differ for any biological or technical reasons. In response to these needs, a number of normalization methods have been proposed. While there seems to be consensus that gene length and library size should be accounted for, methods differ in the type of data features to normalize against and how to do so.

Previous assessments of RNA-seq normalization methods have usually found that common, widely used methods perform well [2–6]. I show here that this is not the case when challenging normalization tasks are considered, suggesting that previous tests may have been overly simplistic. Some evaluations also lacked a gold standard or an optimal solution (but see [2]), in contrast to the results presented below.

The methods evaluated here fall roughly into three groups. Methods that estimate a scaling factor for normalizing libraries (TMM [1] (used in edgeR [7]), DESeq [8] and upper quartile normalization [9]); methods that normalize by gene length and library size (RPKM/FPKM [10], TPM [11]); and a method that aligns z-transformed libraries according to the shape of their overall distribution (zFPKM [12]).

## Results and Discussion

I evaluated the performance of RNA-seq normalization methods according to two commonly encountered scenarios that present challenges for normalization. In the first scenario, I used RNA-seq data from a developmental time course study of *Xenopus tropicalis* embryos [13] that contains spike-in RNA standards [14]. For each normalization method, I asked how well standard RNA levels compared across the dataset. I used the inverse of the coefficient of variation of standard RNA levels across time as a measure of method precision.

The distribution of normalization precision over 45 RNA standards for the different normalization methods is shown in Figure 1A. Precision values for different standards differed considerably for all methods. Most surprisingly, neither method performed markedly better than un-normalized, raw read counts. Only the median precision of zFPKM and upper quartile normalization were above the 75% quantile of the precision of raw read counts; all other methods’ medians fell within the interquartile range. Although its total precision range was the largest, zFPKM had the highest overall precision (median = 0.73 ±0.11 [sample standard deviation]). Upper quartile was the only normalization method with its entire interquartile range above that of raw read counts, showing a very low variation in precision between RNAs (despite a long tail). Nominally, TMM was the only additional method with a higher median precision than raw read counts. I also note that standard RNAs added at higher concentration levels consistently resulted in higher precision values than those present at low levels (Supplementary Figure S1).

**Figure 1.**
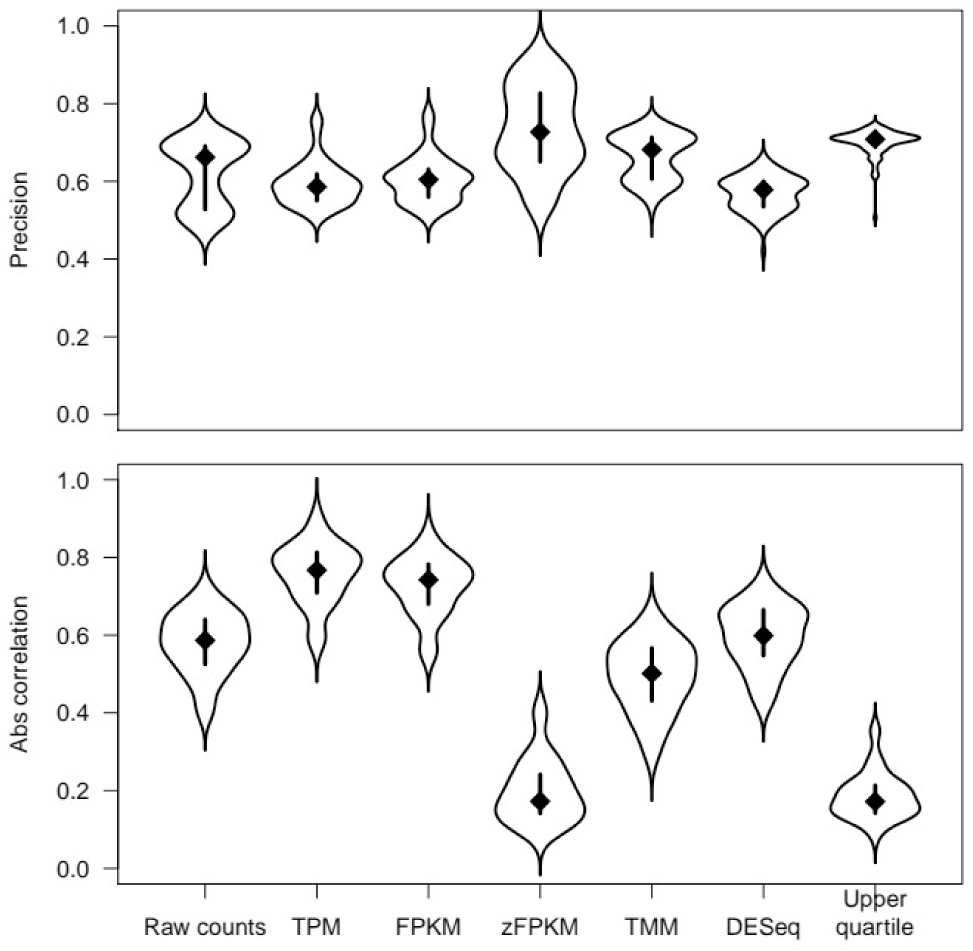
Evaluation of RNA-seq normalization methods using ERCC spike-in RNA measurements across developmental time. (A) Precision was measured as the inverse of the coefficient of variation of RNA levels for 45 spike-ins. Note that spike-ins with high representation in samples tended to have higher precision levels (Supplementary Figure S1). (B) Spearman’s rank correlation between spike-in RNA levels and developmental time. As the developing embryo grows more complex, so does its transcriptome (see [13]). A strong correlation indicates a normalization method’s incapability to account for this fact.

Over developmental time, the transcriptome of the embryo grows ever more complex such that the number of expressed genes increases. Normalization procedures based on the number of genes expressed or transcripts produced may lead to the appearance of stably expressed genes getting more lowly expressed over time while in truth only its relative, but not total abundance decreases. I reasoned that a correlation between stably represented standard RNAs and developmental time would indicate that a normalization method is not able to account for this increasing complexity, a feature that had already been noted in the original study producing the data [13]. I thus calculated Spearman’s rank correlation to estimate this effect. Similar to above, only zFPKM and upper quartile normalization showed reasonably low absolute correlation levels, while all other methods showed correlations of roughly the same strength as raw read counts (Figure 1B). In fact, FPKM and TPM even showed stronger correlations with developmental time than raw read counts.

The theory behind some of the normalization methods provides an explanation for this result. As Wagner and colleagues argue [11], the idea behind many popular normalization methods is to compare methods relative to a so-called relative molar RNA concentration (RMC), the idea that the relative concentration of RNAs is on average constant between samples. In samples from a developing embryo however, this assumption does not hold, as explained above; in turn, RMC decreases over time. Transcriptomes of different complexity may be compared frequently, for instance when comparing distantly related organisms or different tissues. Given the surprising finding that commonly used normalization methods showed a strong bias when faced with such a scenario, it seems wise to argue against the use of these methods under such scenarios.

For the second normalization scenario, I downloaded RNA-seq data from liver tissue from four different species (human, macaque, mouse and chicken) that was generated in three different laboratories. One lab produced data for all four species using 76-bp, single-end sequencing [15]; one group produced data for the three mammalian species using 100-bp, paired-end sequencing [16]; and one group produced data for chicken only, also using 100-bp, paired-end sequencing (Chickspress database [17]).

This scenario presents a different, yet also difficult, challenge as the first scenario. A useful normalization method should be able to normalize libraries such that biological variation is preserved while technical artifacts such as those from experimentation in different labs are removed. Thus, after mapping, I collected 1:1:1:1-orthologous genes, normalized data and performed principal component analysis (PCA) to visualize if biological or technical factors were emphasized after normalization.

In a plot of PC1 and PC2, normalization with three (DESeq, TMM and upper quartile) of the methods resulted in a pattern overtly identical to that of un-normalized, raw read counts (Figure 2). Instead of clustering by phylogeny or sample sex (Supplementary Figure S2), these PCAs show clustering according to source lab. PCA on data normalized with the other three methods (TPM, FPKM, zFPKM) clustered data recognizably according to phylogenetic patterns in the data, albeit to different degree. PCA on data from all three methods split chicken from mammalian samples on PC1, following the deepest phylogenetic split. PC2 showed less resolution regarding further variation, except for data normalized with zFPKM. PCA on the latter split mouse from primates on PC2 (Figure 2) and macaque from human on PC3 (Supplementary Figures S3, S4).

**Figure 2.**
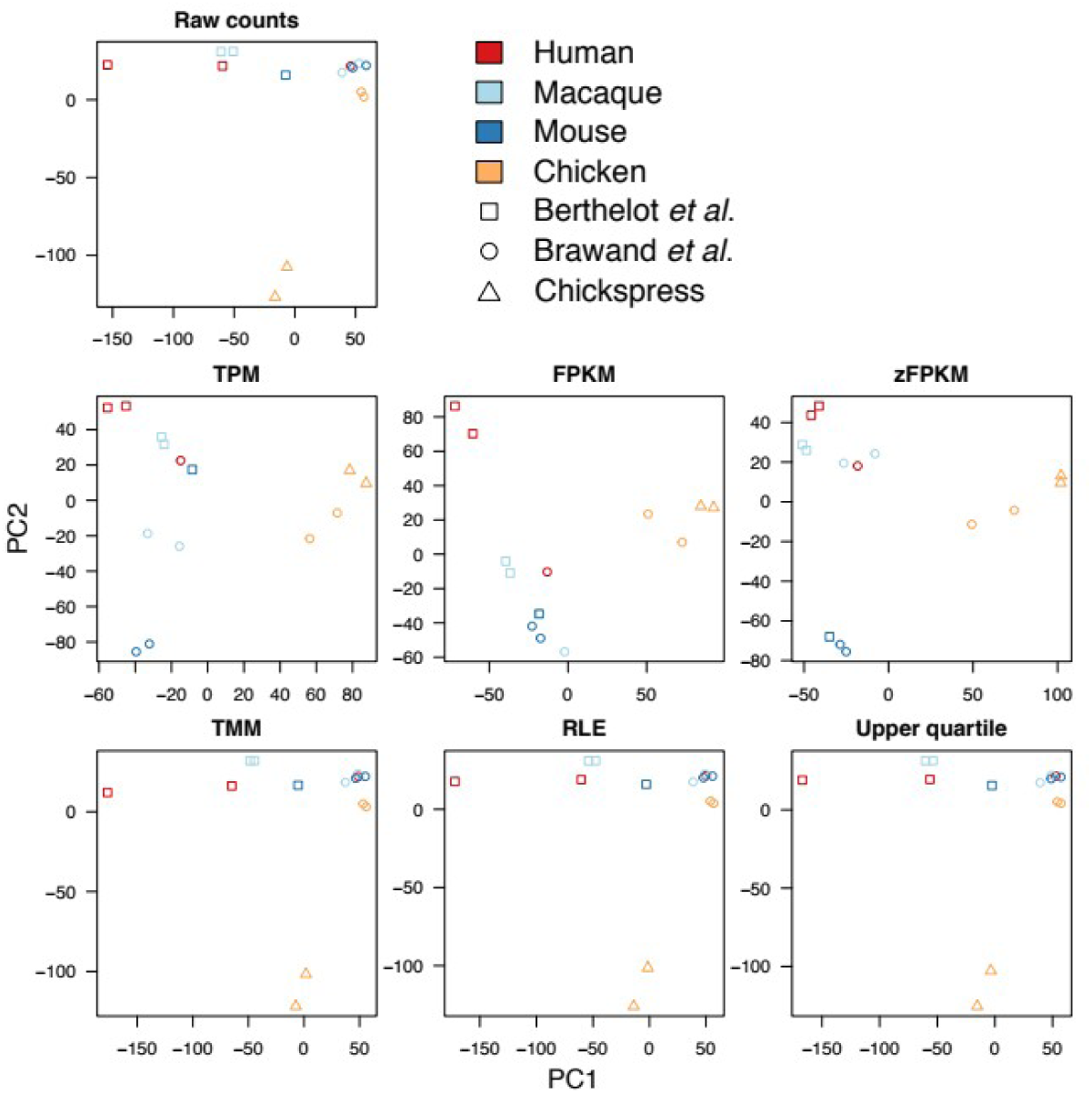
PCAs for un-normalized raw counts, TPM, FPKM, zFPKM, TMM, DESeq, and upper quartile normalizations. Source organism is color coded, source lab is indicated by symbol. See Supplemental Information for PCA with sample sex indicated (Figure S2) and for plots of PC3 (Figures S3 and S4).

The three normalization methods that use scaling factor normalization performed poorly on this task by emphasizing technical artifacts. The inclusion of gene length normalization improves the behavior substantially, but only the method using z-transformation performs arguably optimal. It would certainly be possible (and advisable) to use an approach like z-transformation on data normalized using other methods. Having this feature built-in as a standard speaks for using zFPKM as a go-to method in RNA-seq normalization.

## Summary

In this review, I showed that commonly used RNA-seq normalization methods do not perform well in two challenging, yet common, experimental tasks. This is also true for two normalization procedures that are part of widely used methods for testing differential gene expression, although this may be addressed by downstream computation within the methods. I further show that two methods that are commonly used for comparison and visualization do not perform well with transcriptomes of different complexity. Finally, a rather unknown method, zFPKM [12], performed better than other methods in both normalization scenarios.

## Methods

For scenario 1, I acquired RNA-seq libraries that were sampled at even hours during the first 24 hours (accession numbers below). I used HISAT2 [18] with standard options to map data to the *Xenopus tropicalis* genome RefSeq v. 9.1 with the ERCC reference transcripts [14] added. I used StringTie [19] to summarize mapped reads against RefSeq xenTro v. 9.1 Gnomon gene predictions, including annotations for the ERCC reference transcripts. StringTie reports TPM, FPKM, and raw count values (the latter as “coverage,” which are raw counts normalized by gene length). zFPKM values were calculated from FPKM values according to [12] using an R script. Values for TMM, DESeq, and upper quartile normalization methods were calculated from raw counts using the calcNormFactors() function in the edgeR package [7].

I evaluated normalization methods according to the consistency of ERCC standard RNA levels across time. Standard RNAs that were deemed not expressed by zFPKM (log_2_ zFPKM < −3) were ignored for all methods. First, I calculated precision as the inverse of the coefficient of variation of standard RNA levels across time. Second, I used Spearman’s rank correlation of standard levels against time as a measure for a trend through time.

For scenario 2, I downloaded a number of RNA-seq libraries for liver tissue from four different vertebrates (Supplementary Table S1). I mapped data to the ENSEMBL (v. 92) reference genomes (v. 39) for the respective species and summarized mapped reads against ENSEMBL reference annotations. I used biomaRt [20] to map 1:1:1:1 orthologous genes to human and performed PCA. Statistical analyses were performed using R v. 3.5.0. For availability of scripts, see below.

## Availability of data and material

Sequencing data is available on the SRA under the accessions PRJNA275011, PRJNA143627, PRJEB11781 and PRJNA204941. All scripts are available at https://bitbucket.org/severinevo/normalise/.

## Authors’ contributions

SU designed the study, analyzed data and wrote the manuscript.

## Acknowledgments

I thank J. P. Noonan for comments on an earlier version of this manuscript.

